# Lymphocytic choriomeningitis arenavirus utilises tunnelling nanotube-like intercellular connections for cell-to-cell spread

**DOI:** 10.1101/2023.06.28.546878

**Authors:** Owen Byford, Amelia B. Shaw, Hiu Nam Tse, Alex Moon-Walker, Erica Ollmann Saphire, Sean P. J. Whelan, Martin Stacey, Roger Hewson, Juan Fontana, John N. Barr

## Abstract

The *Arenaviridae* family within the *Bunyavirales* order of segmented RNA viruses contains over 50 species grouped into four genera, *Antennavirus*, *Hartmanivirus*, *Mammarenavirus* and *Reptarenavirus*. Several mammarenaviruses are associated with fatal hemorrhagic fevers, including Lassa, Lujo and Junin viruses. The mammarenavirus member lymphocytic choriomeningitis virus (LCMV) is largely non-pathogenic to humans and represents a tractable model system for studying arenavirus molecular and cellular biology. During infection of cells in culture, a high proportion of LCMV spread is between directly neighbouring cells. Consistent with this observation LCMV-infected cells extrude multiple tunnelling nanotube (TNT)-like structures forming intercellular connections that could provide a route of cell-to-cell spread. To investigate this, we used recombinant LCMV with engineered epitope tags in glycoprotein spike (GP-1) and matrix (Z) proteins, alongside nucleoprotein (NP) antisera, to reveal that all three major structural proteins co-localised within TNT-like connections. Furthermore, utilising fluorescent in situ hybridisation (FISH) we showed NP also co-localised with LCMV genomic sense RNA. Taken together, these observations suggested LCMV virions pass between cells through intercellular connections to infect new cells. Consistent with this, addition of a potent LCMV neutralising antibody to supernatants during infection failed to block LCMV spread through cultures, revealing that cell-to-cell connectivity plays a major role in LCMV transmission. This is the first report of cell-cell infection via TNT-like connections for any species of the 14 families within the *Bunyavirales* order. This study furthers our understanding of how arenaviruses manipulate the host to establish infection, which may aid in the development of effective anti-viral therapeutics.

**IMPORTANCE:** Arenaviruses include some of the most serious human pathogens in existence, although no clinically approved vaccines or therapies are currently available to prevent their associated disease. As with most pathogens, transmission of arenaviruses from one cell to another is a critical aspect of infection and resulting pathogenicity. Here, we showed that model arenavirus lymphocytic choriomeningitis virus (LCMV) can spread between cells without exposure to the extracellular space. We visualized the three major LCMV structural proteins, namely nucleoprotein, glycoprotein spike and matrix co-localized along with genomic RNA within tubular structures connecting adjacent cells. The use of a potent neutralizing antibody to block the extracellular route of LCMV transmission reduced spread within cultured cells to approximately half that of untreated cultures. Taken together, these results suggest intercellular connections represent important conduits for arenavirus spread. This information will aid in the development of antiviral strategies that prevent both intra- and extracellular transmission routes.

## INTRODUCTION

The *Bunyavirales* order of segmented RNA viruses comprises over 500 named viruses divided into 14 families, of which the *Arenaviridae* family currently contains 56 species subdivided into four genera: *Antennavirus*, *Hartmanivirus*, *Mammarenavirus* and *Reptarenavirus* [1]. Mammarenaviruses are designated as Old World (OW) or New World (NW) viruses based on geographical location of their isolation and prevalence, which also correlates with distinctive genetic lineage [2]. Several mammarenaviruses cause serious human disease, such as the OW Lassa virus (LASV) and NW Junín virus (JUNV) [1, 2], both notable for their association with fatal haemorrhagic fevers [3]. Currently, no specific antiviral therapeutics or FDA-approved vaccines exist to target any member of the *Arenaviridae* family [4]. Together, these factors have contributed to LASV, JUNV and other arenaviruses being defined as hazard group 4 pathogens, requiring the highest biosafety level (BSL) 4 containment.

The prototypic species within the *Mammarenavirus* genus is the OW lymphocytic choriomeningitis virus (LCMV) for which rodents are the main vector, with the common house mouse *Mus musculus* acting as primary host [2]. Rodent-to-human LCMV transmission frequently occurs, yet severe disease only arises in rare cases. This is most often in immunocompromised patients or neonates, where infection can result in aseptic meningitis [5, 6], with LCMV described as an underrecognised agent of neurological disease [6]. Specifically, the LCMV Armstrong strain is a hazard group 2 virus, requiring only BSL-2 containment, thus acting as a relevant research model for the more pathogenic species of the *Mammarenavirus* genus, due to similarities in structure and function.

Within mature virions, the viral-associated (vRNA) is coated by the viral nucleoprotein (NP) and interacts with the viral RNA-dependent RNA polymerase (RdRp, L protein) to form the viral ribonucleoproteins (RNPs). These are surrounded by a lipid bilayer, which is lined by the viral matrix (Z) protein and contains protruding viral spikes. All species within the *Mammarenavirus* genus have genomes consisting of two segments, small (S) and large (L), which together encode four structural proteins. All species of this genus express their genes using an ambi-sense strategy; initially, input vRNA S and L segments are transcribed by the viral RdRp, following entry and uncoating, to generate mRNAs that encode the NP and RdRp, respectively. Subsequently, replication of the full-length S and L vRNAs produces S and L anti-genome RNAs (agRNA), which act as templates for synthesis of mRNAs encoding glycoprotein precursor (GPC) and Z proteins [7, 8]. The GPC is post-translationally cleaved to form a trimeric assembly of structural proteins glycoprotein-1 (GP-1), glycoprotein-2 (GP-2) and a stable signal peptide (SSP) [9].

Although LCMV has been intensively studied from an immunological perspective, many molecular aspects of its replication cycle remain poorly understood. One such area is the dependence of viral multiplication on host cell components. While the catalogue of critical cellular factors is increasing, with recent additions to the list including components of the coat protein complex I (COPI) and adaptor protein complex 4 (AP-4) [10] the chaperonin TCP-1 Ring Complex TRiC/CCT [11] and sialomucin core protein 24 (CD164) as a secondary receptor for entry [12], many gaps in our knowledge remain.

Here, we investigated the ability of LCMV to infect cells via intercellular connections rather than the canonical extracellular infection route that involves egress and subsequent entry. Many viruses, including coronaviruses [13], retroviruses [14–17], alphaviruses [18] and orthomyxoviruses [19], have all been shown to utilise intercellular connections as a method of cell-cell spread. Possibly this mechanism represents an efficient means of transmission that does not require intact virion formation and allowing evasion of the host immune response [20]. One such class of connections are tunnelling nanotubes (TNTs), which are membranous tubular structures possessing a backbone rich in filamentous-actin (F-actin) that vary in diameter between 50 to 200 nm and can extend for distances of up to 100 µm [21]. TNTs allow cellular connectivity and have been shown to permit the transfer of varied components between distant cells, including organelles, vesicles, signalling molecules, and ions [17].

Using recombinant LCMV variants with engineered epitope tags, we showed that the three major LCMV structural proteins, NP, GP-1 and Z, all localised within TNT-like structures during infection. Furthermore, fluorescent in situ hybridisation (FISH) showed these components also co-localised with LCMV genomic sense RNA. Taken together, these observations suggest intact LCMV virions and potentially also RNPs may pass between cells through these connections, and the finding that LCMV infection progressed in the presence of a potent neutralizing antibody suggested that movement of viral components through such connections represents an efficient route of cell-cell transmission.

This is the first report of cell-cell infection via TNT-like connections for any species within the *Bunyavirales* order. This study furthers our understanding of how arenaviruses manipulate the host to establish infection, which may aid in the development of effective antiviral therapies.

## RESULTS

### Observation of rLCMV-EGFP infection of cultured cells suggests transmission may involve direct cell-to-cell connectivity

During routine LCMV infections of cultured cells, when omitting semi-solid media overlay, we noted that infected cells formed discrete foci rather than being randomly dispersed throughout the culture. To further examine LCMV spread, cells were seeded and infected with previously described recombinant LCMV expressing eGFP (rLCMV-EGFP) [22] at an MOI 0.001, without semi-solid media overlay. Whole well live-cell analysis was then used to enable continuous monitoring of infected cell clusters every 6 hours post infection (hpi) (Figure 1). At 6 hpi, eGFP expression was first visualised within a single infected cell (Figure 1; white arrow). As infection progressed with time, an increasing number of neighbouring cells became infected, culminating at 42 hpi when a discrete fluorescent focus was visualised, with few cells outside of this showing evidence of infection. This finding suggested that LCMV spread within a culture may involve transmission via direct cell-to-cell contact.

**Figure 1.**
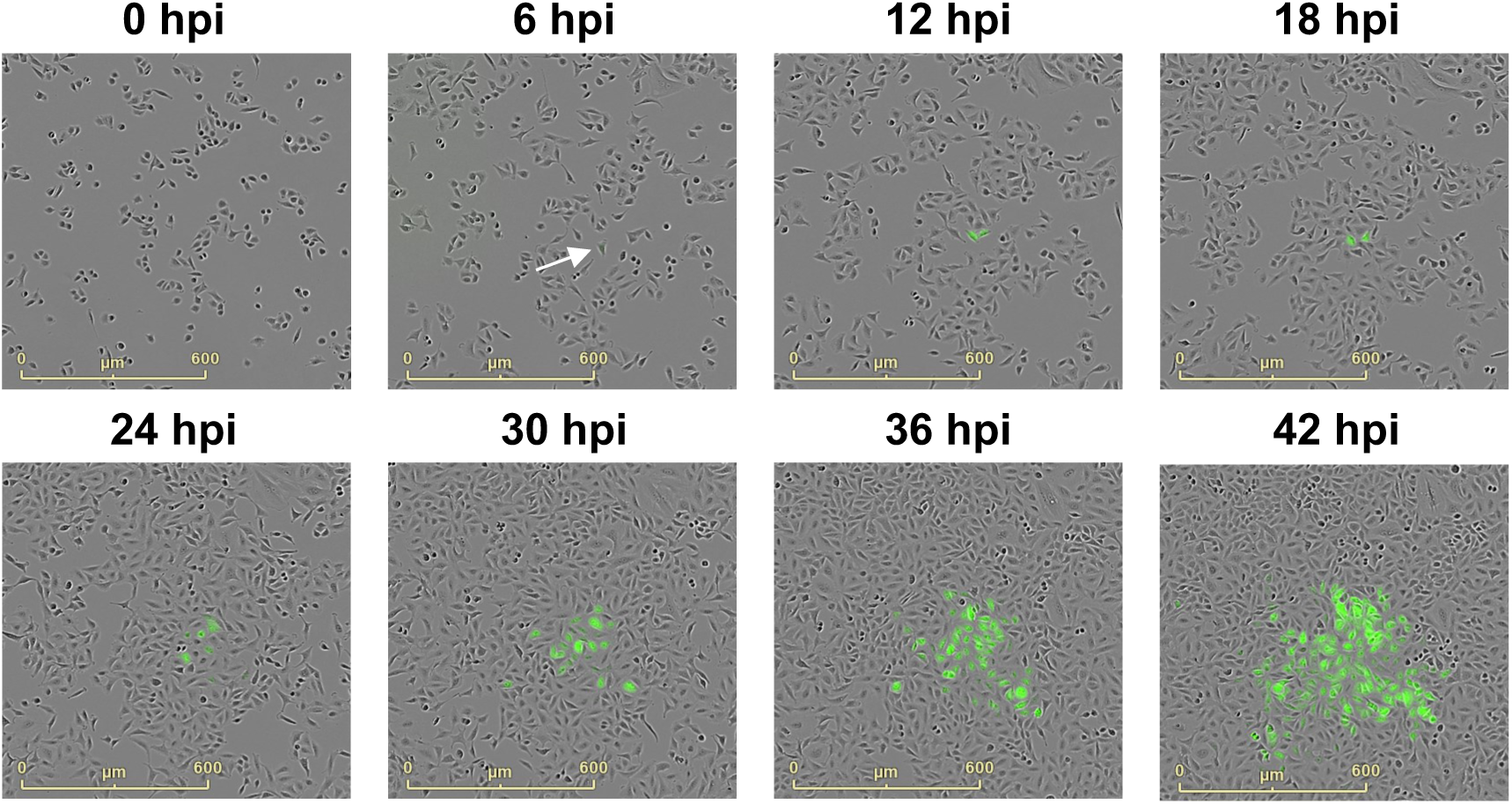
Spread of LCMV infection within a culture is largely confined to adjacent cells leading to focus formation. A549 cells were seeded and simultaneously infected with rLCMV-eGFP at multiplicity of infection (MOI) of 0.001, with cells cultured in liquid medium. Phase contrast and eGFP fluorescence whole well images were taken every 6 hours, and subsets of cells monitored until 42 hpi. A single infected cell (white arrow) was monitored for the duration of the experiment.

### LCMV infected cultures contain inter-cellular connections that contain viral components

To further investigate the role of cell-to-cell contacts in LCMV spread, we first needed to detect such connections, and confirm that viral components could be detected within. To do this, A549 cells were infected with our previously described rLCMV-GP1-FLAG [10] and fixed at 24 hpi with staining to detect the cellular distribution of the LCMV GP-1 spike alongside F-actin, used to aid in the demarcation of the cell periphery and visualization of intercellular connections. Cells were visualized using stimulated emission depletion (STED) microscopy, which revealed abundant intercellular connections between cells comprising a complex F-actin architecture, with multiple F-actin bundles running parallel with the long axis of the tube, both bordering the tube as well as forming an internal central core.

In the image shown, one prominent F-actin filament emanated from within the cell on the right of the image and extend through the connection (Figure 2A, arrows). The staining of GP-1 within the two interconnected cells was different; in the right-hand cell, GP-1 was widely-distributed with abundant intense signal suggestive of a late-stage infection, whereas in the left–hand cell, the GP-1 signal was of low intensity, suggestive of an early-stage infection. Within the cell-cell connection, the GP-1 signal exhibited a gradient of abundance, which decreased with distance away from the late-stage infected cell. The majority of GP-1 was located proximal to, but not coincident with, the central F-actin fibres (Figure 2A, arrows). This observation was corroborated with line scan analysis (Figure 2B; corresponding to dotted line in panel A [start 1, end 2]), which showed the peak signals corresponding to F-actin did not precisely overlap with GP-1, but instead were separated by approximately 90 nm. Interestingly, super resolution measurement of regions within the cell-cell connection confirmed GP-1 puncta were of a size (100-150 nm) expected for intact virions. Taken together, these findings suggest that intercellular connections may act as conduits for the transport of LCMV components, or even intact virions, between neighbouring cells.

**Figure 2.**
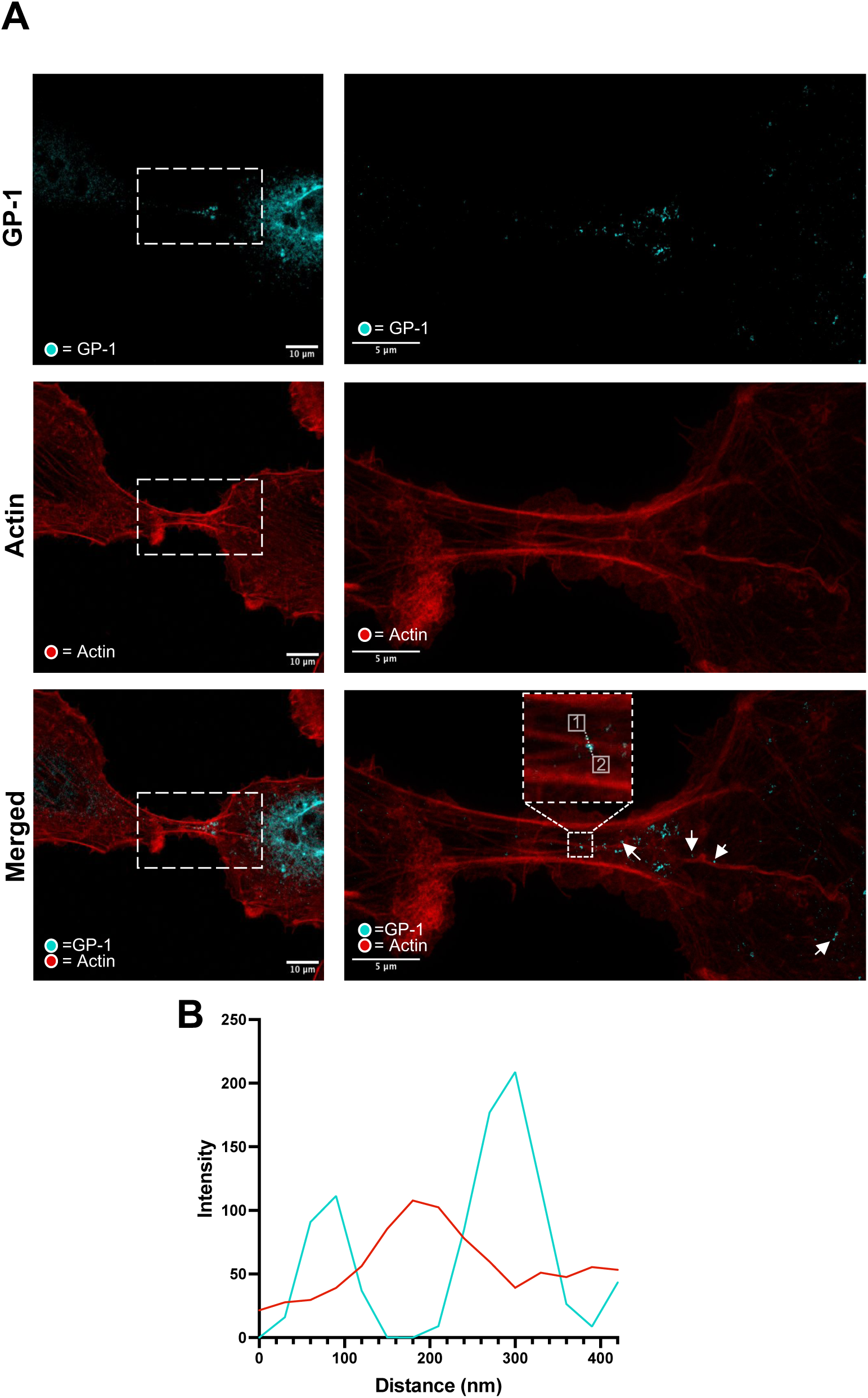
A549 cells infected with rLCMV-GP1-FLAG form cell-cell connections that contain F-actin and virion components. A) A549 cells were infected with rLCMV-GP1-FLAG at an MOI of 0.2, and at 24 hpi, cells were fixed, stained for GP-1-FLAG (cyan) and F-actin (phalloidin, red), then visualized by stimulated emission depletion microscopy (STED). The left panels show two connected cells, with a white dashed box indicating the area of the zoomed images shown as the right panels, with a further zoomed image also shown where line scan analysis was performed. Punctate regions of LCMV GP-1 positioned along an F-actin branch are indicated by arrows. (B) Line scan analysis of the region of interest in the zoomed merged image (white box), showing GP-1 peak intensities do not closely correspond to those of F-actin at this resolution.

### Cell-cell connections contain both actin and tubulin

To further characterise the cellular components involved in forming cell-cell connections, A549 cells were infected with rLCMV-GP1-FLAG for 24 hpi. At that time, cells were stained for the cytoskeletal components F-actin (red) and β-tubulin (green) alongside GP-1 (cyan), followed by visualisation using indirect immunofluorescence (IF) confocal microscopy.

Consistent with our findings above (Figure 2), at 24 hpi GP-1 was predominantly localised within perinuclear regions, but also within puncta throughout the cytosol and on the cytosolic face of the plasma membrane (Figure 3). As expected, inspection of the intercellular spaces revealed GP-1 was also present within filamentous projections that extended between adjacent cells (Figure 3). In addition, IF analysis revealed that both F-actin and β-tubulin were present within the tubular cell-cell connections. These cell-cell connections varied in size but often appeared to connect an infected cell containing abundant GP-1 to cells devoid of GP-1, likely uninfected. Furthermore, in many cases, multiple GP-1 puncta were observed extending beyond cell-cell connections into the uninfected cell (Figure 3; arrows), consistent with the connections being open-ended. As these intercellular connections share some of, but not all, the distinguishing characteristics of TNTs, namely rich in F-actin and open-endedness, hereafter we refer to these connections as ‘TNT-like’.

**Figure 3.**
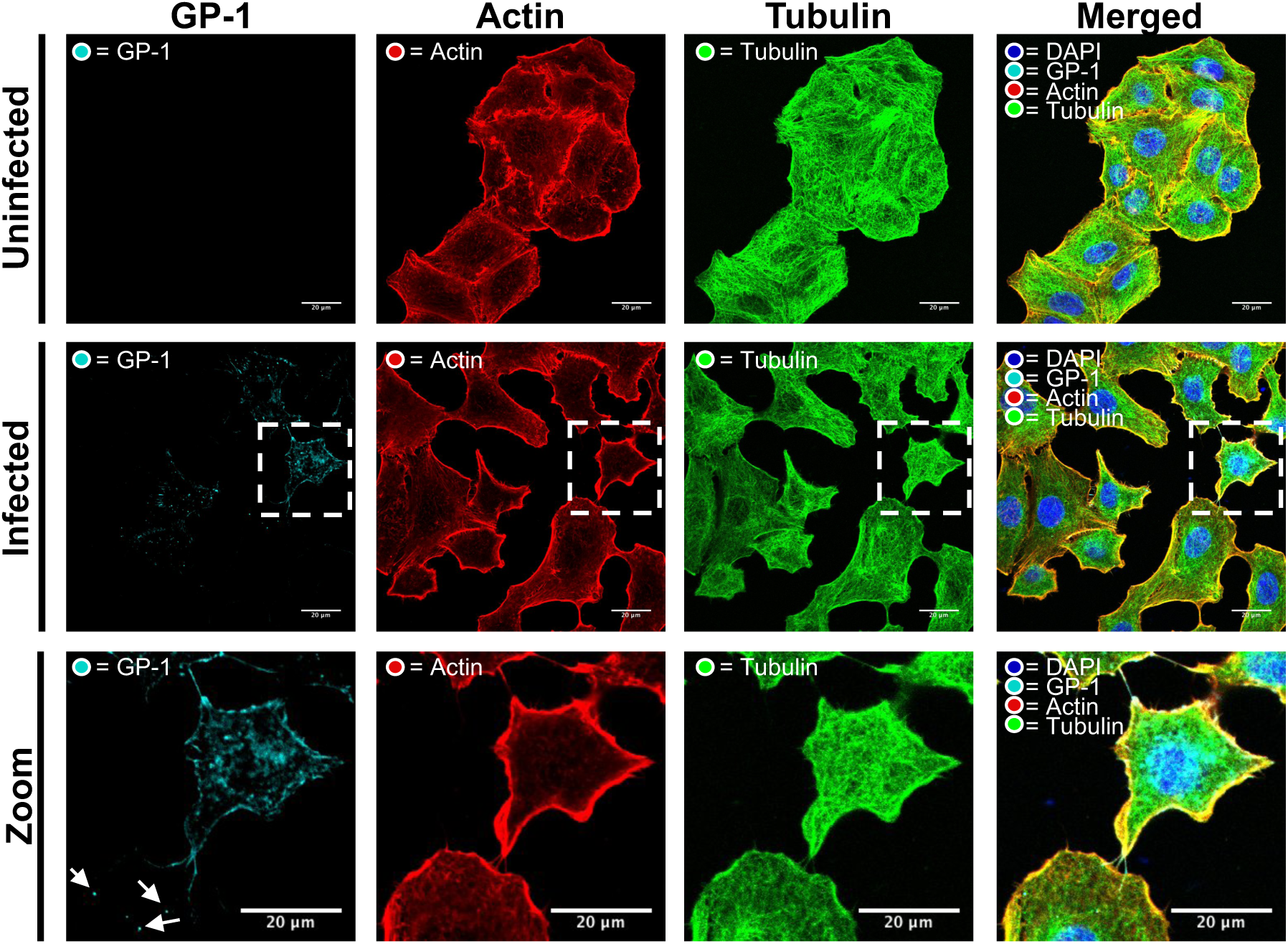
Intercellular connections between LCMV-GP-1-FLAG infected cells contain both F-actin and tubulin. A549 cells were infected with rLCMV-GP1-FLAG at an MOI of 0.2. At 24 hpi, cells were fixed and then stained with DAPI and phalloidin (actin) alongside antisera specific for GP-1 (cyan) and a β-tubulin-specific Affimer (green), and then visualized by confocal microscopy. A zoomed-in region of interest (white box in the central row) is shown on the bottom row, where multiple cell-cell connections are visualized by staining with both tubulin and F-actin. Additional points of interest are indicated by arrows to highlight punctate regions of LCMV GP-1 present in neighbouring cells, extending beyond F-actin and β-tubulin-stained tubular structures.

### Rescue of a recombinant LCMV variant with GP-1 FLAG tag alongside Z HA tag (rLCMV-GP1-FLAG-Z-HA)

To investigate the involvement of LCMV NP, Z and GP-1 within TNT-like connections, we collated two previously described infectious rLCMV variants, namely rLCMV-Z-HA [22] and rLCMV-GP1-FLAG [10], generating the rLCMV reassortant rLCMV-GP1-FLAG-Z-HA (Figure 4A). For this virus, the S segment expressed FLAG-tagged GP-1 and the L segment expressed a HA-tagged Z, allowing simultaneous detection of LCMV NP (using NP antisera), Z and GP-1, the three major structural components of the virion, within infected cells (Figure 4B). Western blot analysis confirmed successful rescue of rLCMV-GP1-FLAG-Z-HA (Figure 4C), with the presence of LCMV NP at 5 days post transfection indicating successful recovery of rLCMV-GP1-FLAG-Z-HA. Subsequently, supernatants were transferred to fresh BHK cells and harvested at 2 days post infection (dpi). Western blot analysis revealed successful recovery of rLCMV-GP1-FLAG-Z-HA, and the relative NP abundance for all mutants recovered were comparable to WT (Figure 4C). Subsequently, rLCMV-GP1-FLAG-Z-HA stocks were amplified and titrated in BHK cells, reaching a titre of 5 x 10^5^.

**Figure 4.**
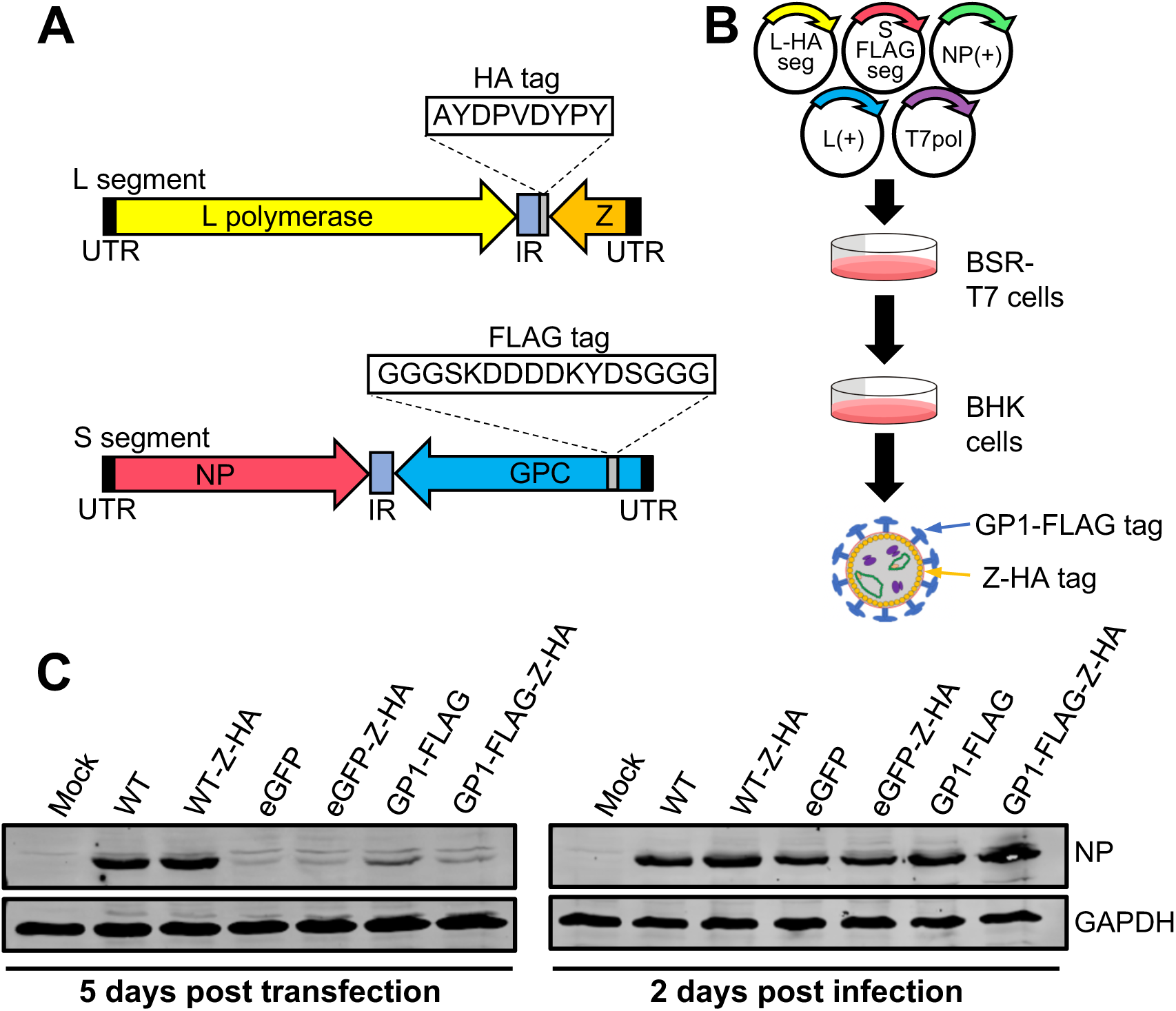
Rescue of a recombinant LCMV variant with GP-1 FLAG tag alongside Z HA tag (rLCMV-GP1-FLAG-Z-HA). (A) Schematic of LCMV S and L segments with respective tag insertions. (B) BSR-T7 cells were transfected with cDNAs expressing a LCMV S segment encoding a FLAG tagged GP-1 (S-FLAG segment), an L segment encoding a HA tagged Z (L-HA segment), as well as NP and L open reading frames as support plasmids, L(+) and NP(+) and a plasmid expressing T7 RNA polymerase, T7pol. At 5 days post transfection (5 dpt), cellular supernatants were transferred to BHK cells, and at 2 days post infection (2 dpi), cell lysates were analysed by western blotting using NP antisera. (C) Western blot analysis of transfected BSR-T7 and infected BHK-21 cell cultures confirmed rLCMV-GP1-FLAG-Z-HA rescue (alongside all other mutants), using antisera specific for LCMV NP and GAPDH as loading control.

### LCMV GP-1, NP and Z puncta co-localise within TNTs during infection

Next, rLCMV-GP1-FLAG-Z-HA was used to assess the localisation of NP, Z and GP-1 within infected cells and TNT-like connections. A549 cells were infected with rLCMV-GP1-FLAG-Z-HA at a MOI of 0.25, and the cellular localisation of NP (red), GP-1 (cyan) and Z (green) was assessed using specific antisera via widefield IF microscopy (Figure 5).

**Figure 5.**
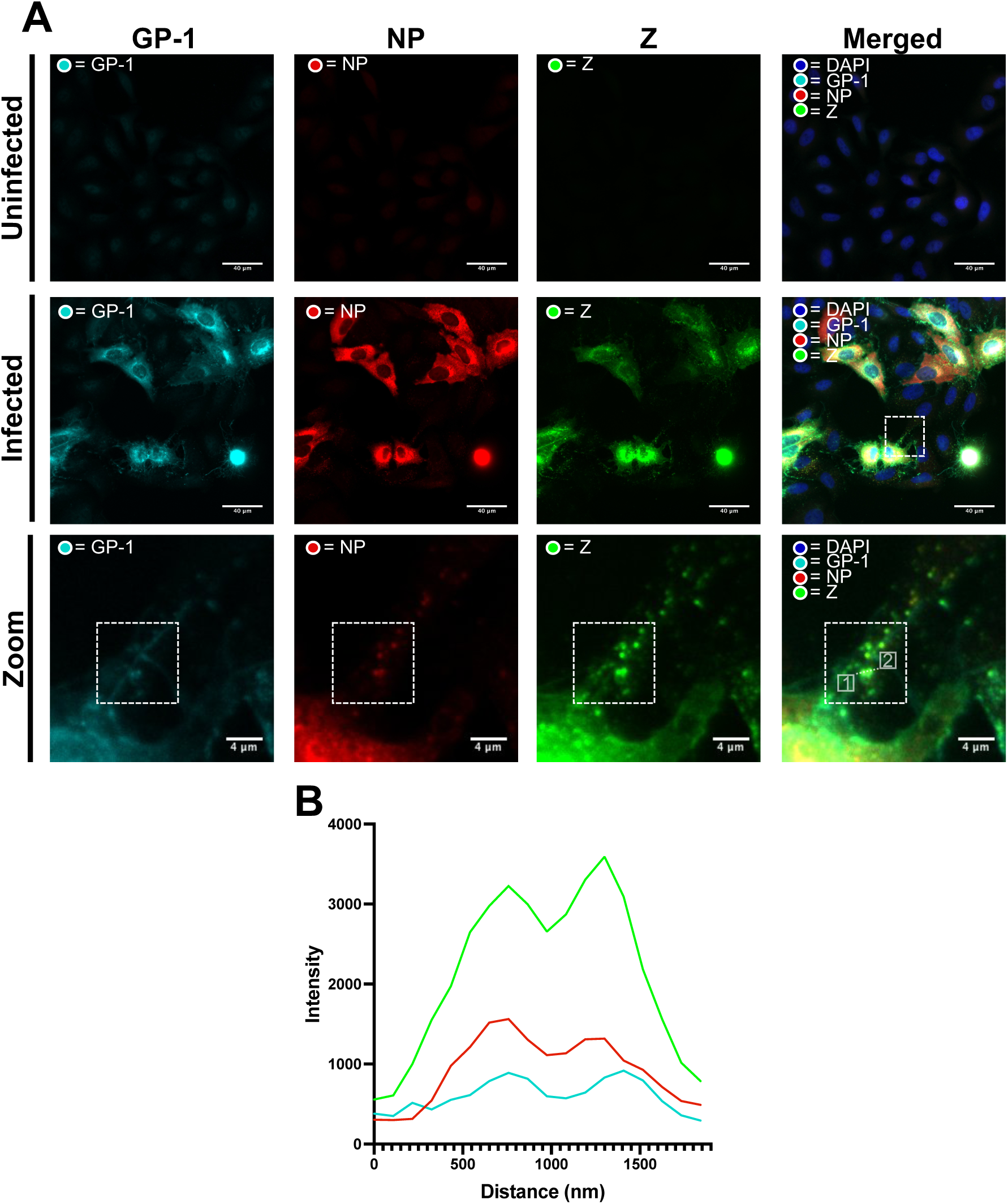
GP-1, NP and Z puncta co-localise within TNTs during rLCMV-GP1-FLAG-Z-HA infection. (A) A549 cells were infected with rLCMV-GP1-FLAG-Z-HA at an MOI of 0.25. At 24 hpi, cells were fixed, stained for GP-1 (cyan), NP (red) and Z (green) and visualized by widefield microscopy. LCMV GP-1, NP and Z staining are distinct and present within TNTs as small puncta (∼200 nm). Uninfected cells and infected cells are shown. A zoomed-in region of interest (white box in the central row) is shown on the bottom row. (B) Line scan analysis of the region of interest in the zoomed merged image (dashed line in the bottom right panel of A), revealed that although the signal intensities vary, the intensity peaks for NP, GP-1 and Z are closely aligned.

IF analysis at 24 hpi revealed the distribution of each of NP, GP-1 and Z was distinctive. In agreement with our previous findings, LCMV NP formed discrete perinuclear puncta, as well as being widespread throughout the cytoplasm [22]. LCMV GP-1 localisation differed to that of NP, being predominantly punctate and perinuclear as well as being abundantly distributed along the plasma membrane. Interestingly, Z showed a similar cellular distribution to GP-1, with clear and abundant Z/GP-1 co-localisation in perinuclear regions, and co-localization at discrete locations on the inner face of the plasma membrane, possibly representing virus assembly sites.

Within TNT-like connections between LCMV infected and uninfected cells (Figure 5A; white boxes in central row and bottom row) multiple puncta were identified that contained NP, GP-1 and Z. Due to their discrete spherical shape and the presence of all three major structural proteins, these objects likely represented assembled virions. In further support of this, line scan analysis across two discrete puncta (at 650 – 867 nm and 1300 – 1408 nm) confirmed the peak intensities for each viral component were spatially-aligned (Figure 5B, corresponding to dotted line in panel A, bottom row [start 1, end 2]).

### LCMV NP and viral genomes co-localize within TNT-like connections

The detection of LCMV NP, GP-1 and Z co-stained puncta inside TNT-like connections suggested these objects represented assembled LCMV virions. To further investigate this possibility, we performed fluorescent *in situ* hybridisation (FISH) using probes designed to detect negative sense LCMV vRNA genomes, which would be predicted to also co-locate to assembled virions.

To achieve this, we generated a set of fluorescently labelled probes specific for the NP coding region of the negative sense rLCMV S RNA segment, with which we performed FISH on A549 cells infected with rLCMV at an MOI of 1.0 at 24 hpi using wide-field microscopy. To specifically detect assembled LCMV RNPs, which represent the form in which vRNAs are packaged within virions, cells were also co-stained with antisera specific for LCMV NP.

As anticipated for an MOI of 1.0 at this time point, most cells appeared infected as evidenced by staining by FISH and NP (Figure 6). Within cells, NP distribution was consistent with previous findings, and FISH analysis revealed close co-localisation of NP and LCMV S segment vRNA at several locations within the infected cells (Figure 6A, central row) sometimes in perinuclear locations, but also as dense puncta throughout the cytosol. Within TNT-like connections, signals corresponding to NP and S segment vRNA were both visualised (Figure 6A, white boxes in central row and bottom row). While staining corresponding to both NP and S segment vRNA was weak within TNT-like filaments, discrete punctate regions were apparent, within which NP and RNA signals closely co-localized. This was corroborated using line scan analysis (arrow), which revealed peaks corresponding to both NP and S vRNA were closely coincident (Figure 6B; corresponding to dotted line in panel A, bottom row [start 1, end 2]). The proximity of the LCMV NP and S segment vRNA signals was consistent with their location within the assembled RNP, with the small offset in peak spatial intensity likely a reflection of the differing approaches utilised for target detection; the FISH probes bind directly to the S vRNA target and emit signal from fluorogenic probe-bound dyes. In contrast, the NP signal results from binding of primary and fluorescent secondary antibodies, thus some spatial separation between target and fluorophore was expected.

**Figure 6.**
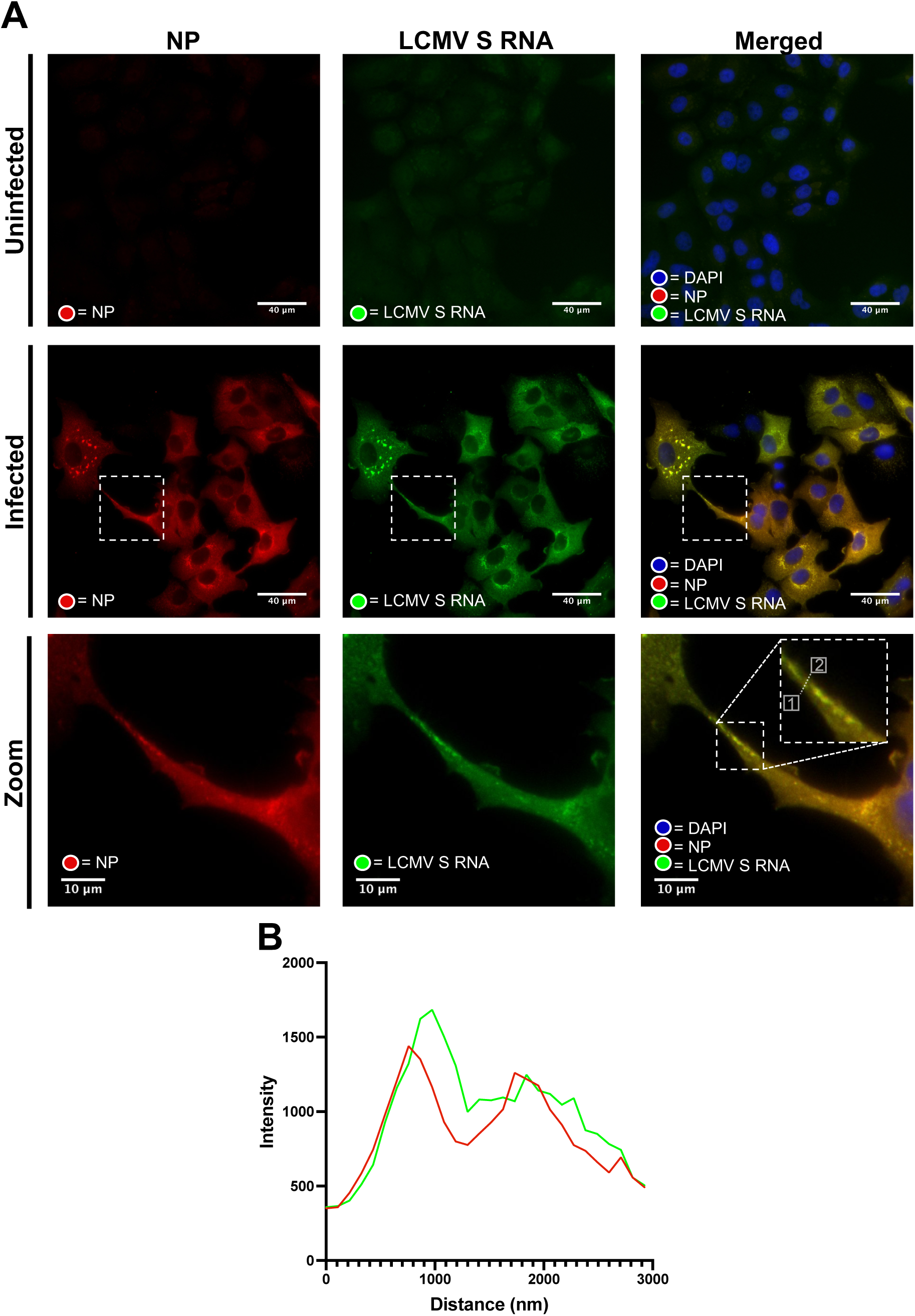
LCMV vRNA is present alongside NP in TNT-like connections. (A) A549 cells were infected with rLCMV at an MOI of 1. At 24 hpi, cells were fixed and then stained for NP (red). Subsequently, fixed cells were incubated overnight with 48 specific FISH probes designed to hybridize with LCMV S vRNA (green). Widefield microscopy was then utilised to visualise NP and S vRNA within TNT-like cellular connections. Cell nuclei were DAPI stained. Uninfected cells and infected cells are shown. A zoomed-in region of interest (white box in the central row) is shown on the bottom row. (B) Line scan analysis of the region of interest in the zoomed merged image (dashed line in the bottom right panel of A), reveals NP and S vRNA peak intensities are closely aligned.

Taken together, this FISH analysis shows that LCMV RNP components are able to travel within TNT-like connections between cells. Given that LCMV NP co-localised with GP-1 and Z within TNT-like connections, it is possible that the co-localised NP and S segment vRNAs puncta also represent assembled LCMV virions. However, we cannot rule out the possibility that assembled RNPs, devoid of both spike and Z matrix proteins, are independently able to travel between cells through TNT-like connections.

### Blocking extracellular LCMV transmission with a potent neutralising antibody reveals that LCMV utilises cell-cell spread during infection

To further confirm that LCMV utilises TNT-like connections for cell-cell transmission, we made use of a recently described LCMV neutralising antibody (M28) that targets GP-1 [23]. We reasoned that when added to infected cell supernatants, this antibody would block LCMV infection via the extracellular route but would not hinder cell-cell transmission via intercellular connections (Figure 7A).

**Figure 7.**
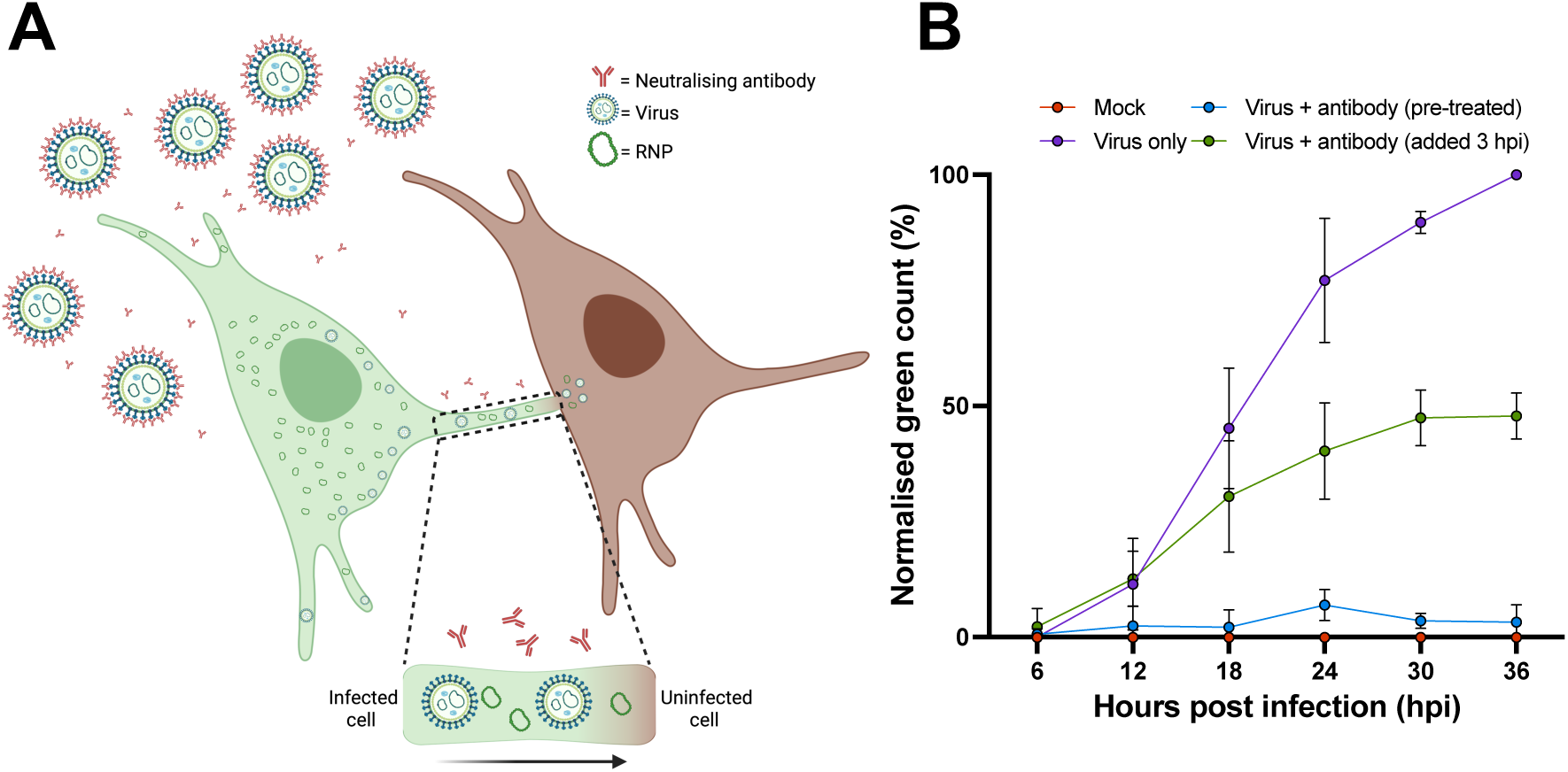
LCMV can efficiently spread through a culture despite blocking extracellular transmission using a potent neutralising antibody. (A). Schematic to illustrate the experimental approach. (B). A549 cells were infected with rLCMV-EGFP at an MOI of 0.1 that were either pre-treated with antibody (blue plotted points; 5 µg/mL antibody for 1 h) or virus only (in the absence of antibody; purple plotted points). For virus + antibody (green plotted points), 5 µg/mL antibody was added to cultures at 3 hpi, following LCMV entry and uncoating, which remained for the duration of the experiment. The total green count (number of green cells) was measured every 6 h, and normalised against virus only (purple) at 36 hpi. The average of three independent experimental repeats is shown (n=3), with error bars showing standard deviation at each time point.

First, to confirm neutralising antibody potency under the selected experimental conditions, M28 neutralising antibody at 5 µg/mL was pre-incubated with rLCMV-EGFP prior to infection of A549 cultures. To measure virus transmission throughout the culture the total green cell count (TGC) was determined at successive 6 hpi time points up to 36 hpi, at which time a TCG value of just 3% was recorded (Figure 7B blue plotted points) and normalized to that of the virus only (in the absence of antibody) control (Figure 7B, purple plotted points). This was statistically indistinguishable from the TCG of mock infected cultures (Figure 7B, red plotted points) and thus confirmed that the antibody effectively blocked extracellular transmission in the culture to background levels of detection.

Next, to investigate whether rLCMV-EGFP could spread within the culture when extracellular transmission was blocked, cultures were infected with rLCMV-EGFP at an MOI of 0.1, with M28 added at 3 hpi, at which time virus entry is complete but virion assembly has not yet commenced [22].This protocol would allow virus to enter these initial cells, but the presence of antibody in the media thereafter would block any subsequent transmission by the extracellular route. These experimental conditions resulted in a TCG value of 47% at 36 hpi (Figure 7B, green plotted points) when normalized to the virus only control, thus revealing significant transmission had occurred throughout the culture despite the continued presence of potent neutralizing antibody.

These results show that LCMV virions can efficiently travel between cells within a culture using a means other than canonical extracellular transmission. Together with the results described above, in which we find evidence of NP, Z and GP-1 within the TNT-like connections, these findings are consistent with TNT-like connections providing an alternative route of infection, allowing cell-to-cell infection without virions being secreted to the extracellular space.

## DISCUSSION

Our previous work has shown that rLCMV-GP1-FLAG [10] and rLCMV-HA-Z [22] represent useful tools for identification of the sub-cellular localisation of LCMV components during multiple stages of the replication cycle. Here, we extended this work to generate an infectious LCMV variant simultaneously expressing both FLAG-tagged GP-1 and HA-tagged Z (rLCMV-GP1-FLAG-Z-HA) to investigate the localisation of GP-1, Z and NP within individual cells.

We showed LCMV infected cells generated TNT-like structures containing both F-actin and β-tubulin connecting infected and non-infected cells. To investigate the role of these connections during infection, we visualized cells infected with rLCMV-GP1-FLAG-Z-HA, where we observed small punctate objects inside TNT-like structures, within which GP-1, Z and NP all closely co-localized, consistent with these objects representing assembled virions.

Furthermore, FISH analysis showed these TNT-like connections contained LCMV S genomic vRNA in close proximity to NP, indicative of RNP structures that would be expected to be found within infectious virions. Thus, taken together, these findings show that the major structural components of LCMV, namely NP, vRNA, Z and GP1 colocalize as discrete virion-sized puncta within TNT-like connections. Consequently, it is difficult to escape the conclusion that these objects are virus particles, raising the possibility that movement of virions through these cellular connections may represent a mode of transmission that does not involve release of virions into extracellular spaces.

In support of this, we showed LCMV could transmit within a culture in the presence of a potent neutralising antibody, which effectively blocked extracellular transmission, revealing that cell-cell LCMV transmission may represent up to 50% of total spread during infection. Taken together, our data suggests that infectious LCMV virions can traffic between cells through intercellular connections, and this mode of virus spread represents a significant proportion of total virus transmission during infection.

This work is the first report describing the utilisation of cell-cell transmission for any species of the *Bunyavirales*. Nevertheless, many other viruses have recently been revealed to rely on TNT or TNT-like structures during infection including species of retroviruses, herpesviruses, alphaviruses, pneumoviruses, orthomyxoviruses, paramyxoviruses, flaviviruses and picornaviurses [24].

Within the *Coronaviridae* family, SARS-CoV-2 virions have recently been visualised to ‘surf’ on the outside of TNT connecting cells during infection, along with evidence to suggest mature virions are also present within the interior of the same structures [13]. This study demonstrated transmission of SARS-CoV-2 between permissive and non-permissive neuronal cells via TNT connections. Thus, TNT connections could potentially act as a mechanism for widespread dissemination of viruses within a host, circumventing different cellular or tissue-type restrictions, such as the requirement for specific entry receptors, allowing spread to tissues and organs distinct from the initially infected target cells.

The reliance of SARS-CoV-2 on TNTs was further investigated with the use of neutralising antibody to block extracellular transmission of a pseudotyped lentivirus, resulting in no significant decrease in cell-cell spread [25]. Thus, it was proposed that cell-cell TNT connections may represent an efficient mode of transmission, allowing evasion of host immune cell responses. Here, we utilised a neutralising antibody to demonstrate that up to 50% of LCMV transmission within a culture result from cell-cell TNT-like connections. It will be interesting to test whether LCMV also exploits TNT-like structures during infection of target host tissues, and also whether this mode of transmission occurs during infection of intact organisms.

Interestingly, influenza A virus (IAV), which like LCMV is an enveloped negative-sense segmented RNA virus, has been shown to utilise TNT connections for spread during infection. However, for IAV the infectious entity is not proposed to be a fully assembled mature virion, but instead is believed to comprise solely the RNP, devoid of both matrix and the spike-embedded envelope [19, 26]. Rab11a is known to be involved in transport of RNPs to assembly sites at the plasma membrane for the more-conventional mode of transmission involving extracellular release, and Rab11 was also shown to play a role in movement of IAV vRNPs across TNT connections [19]. Interestingly, analysis of IAV infection outcome revealed the movement of IAV genomes through TNT connections acted as an important factor to facilitate genome reassortment during co-infections, with important consequences for virus evolution and possible emergence of pathogenic variants [19].

Here, our data is consistent with intact virions being trafficked within TNT-like connections, since we observe GP1 together with Z and NP. However, the results of the FISH analysis also allow the possibility that non-enveloped RNPs are also able to pass through these connections. The distinction between these two possibilities has ramifications for infection of the connected cell. If the transmitting object is an RNP, then presumably this would be able to initiate infection in the same way as if just released from an endocytic vesicle. In contrast, if the transmitting object is an intact enveloped virion, it must be capable of disassembly by fusion with an endomembrane, yet still permit release of the RNP within the cytosol, and how this could be achieved is unclear.

Further investigations are required to understand if the movement of virus components within TNT-like connections is an active or passive process, or if there is a reliance on host cell components for this transport, such as Rab11, as described for IAV, above. Interestingly, STED analysis of rLCMV infected cells stained for GP-1 and F-actin (Fig. 1B and C) showed the corresponding signals were proximal but not coincident. This observation raises the possibility that F-actin associated proteins such as the resident myosin motors may play a role. The host cell motor protein myosin II has been implicated in ‘viral surfing’ for several viruses including murine leukemia virus, avian sarcoma leukosis virus and vesicular stomatitis virus [27]. In the context of HIV infection, host cell motor protein nonmuscle myosin II (NMMII), has been implicated in the movement of Gag and Env through cell-cell TNTs during infection [16]. It is unclear if LCMV requires the involvement of any cellular motors, or if an association with growing F-actin fibres is sufficient for TNT trafficking, and experiments to investigate this are underway.

Another group of viruses that utilize cell-cell spread are the flaviviruses, for which the role of cell-cell transmission of hepatitis C virus (HCV) for evasion of neutralising antibodies is well understood [28]. Direct acting antivirals (DAAs) are effective therapeutics for HCV, yet resistance can occur. Within a recent study, cell-cell transmission was shown to be the primary route of HCV DAA-resistance, leading to infection persistence. By targeting cell-cell transmission alongside DAA treatment, resistance was overcome, and HCV was again eliminated in a cell culture model [28] Overall, this study is the first to reveal a role for TNT-like structures in cell-cell spread during infection for any species of the *Bunyavirales* order. We provide evidence consistent with the trafficking of assembled virions and RNPs with these structures and exploit a potent neutralising antibody to propose up to 50% of transmission within cultured cells may occur through an inter-cellular route. It will be interesting to examine whether other arenaviruses from both OW and NW clades utilise TNT-like connections as transmission conduits, and also to test whether these connections are formed in the context of intact tissues with an infected host.

## MATERIALS AND METHODS

### Plasmid design and virus rescue

Construction of plasmids expressing S and L segments for rescue of rLCMV-WT, rLCMV-eGFP, LCMV-Z-HA [22] and LCMV-GP1-FLAG [10] have been previously described, along with corresponding rescue protocols. Rescue of rLCMV-GP1-FLAG-Z-HA was achieved using these same protocols, by transfection of BSR-T7 cells with cDNAs expressing the S segment of rLCMV-GP1-FLAG along with the L segment of rLCMV-Z-HA (Figure 4). Also transfected were supporting plasmids expressing the ORFs of LCMV NP and LCMV L proteins, as well as bacteriophage T7 RNA polymerase. At 5 days post transfection (dpt), cell supernatants were transferred to BHK cells allowing amplification of any rescued viruses. At 2 days post infection (dpi), viral supernatants were collected and subject to focus forming assays (FFA). Titred viral stocks gained were utilised for bulking in BHK cells at an MOI of 0.001, and subsequently harvested at 3 dpi.

### Virus infections

To generate virus stocks, T175 flasks of BHK cells seeded at 5×10^6^ the day prior were infected at an MOI 0.001. At 3 days post infection, viral supernatant was harvested and centrifuged to remove cell debris (x4000 g), aliquoted (80 µL) and frozen for subsequent viral titration. For all other viral infections, cell monolayers were infected with LCMV at the specified MOI in either serum-free (SFM), 2.5 % or 10 % FBS DMEM, depending on cellular requirements, at 37 °C. After 1 h, the inoculum was removed and SFM, fresh 2.5 % or 10 % DMEM was then applied for the duration of the infection. For synchronised infections, LCMV was bound on ice to cells for 1 h. Subsequently, the inoculum was removed, monolayers washed x3 with PBS and SFM, fresh 2.5 % or 10 % DMEM was then applied for the duration of the infection.

### Viral titration

Determination of virus titres was achieved through focus forming assays (FFA). Viral stocks requiring titration were serially diluted in SFM to infect fresh monolayers of BHK cells seeded at 1 x 10^5^ in a 24 well-plate. After infection, medium containing virus was removed and 1 mL 1:1 ratio of 10 % FBS DMEM to 1.6 % methylcellulose was reapplied, and the cells incubated for a further 3 days. For rLCMV-eGFP titration, the Incucyte Zoom S3 live cell imaging system (Sartorius) was used to image whole wells and detect fluorescent rLCMV-eGFP foci, which were then counted for titre determination. For rLCMV-WT titration, cells are fixed using 4 % (vol/vol) paraformaldehyde (PFA) for 15 mins and washed three times with PBS. Cells were then incubated with permeabilisation buffer (0.3 % [vol/vol] Triton X-100, 2 % [wt/vol] FBS in 1 x PBS) for a further 10 mins, and then washed three times with PBS. Following this, cells were incubated with 1 mL blocking buffer (1 % [wt/vol] BSA in PBS) for 1 h. The cells were then incubated for 1 h with 150 µL/well LCMV NP primary antibody (generated in-house, 1:1000 in blocking buffer), and washed three times with PBS. Following this, cells were incubated for 1 h with 594 Alexa Fluor secondary antibody (Life Technologies; 1:500 in blocking buffer), and subsequently washed four times with PBS. The Incucyte Zoom S3 live cell imaging system was then used to image whole wells of the plate to detect red rLCMV-WT focus forming plaques. Plaques were quantified, and virus titres determined. For rLCMV-FLAG titration, the protocol for rLCMV-WT was followed, yet cells were incubated for 1 h with 150 µL/well LCMV NP (in-house, 1:1000 in blocking buffer) and FLAG (Sigma, 1:500 in blocking buffer) primary antibodies. Following this, cells were incubated for 1 h with 488 (FLAG) and 594 (LCMV NP) Alexa Fluor antibodies (Life Technologies; 1:500 in blocking buffer), then washed four times with PBS.

### Analysis of LCMV infection progression

Trypsinised A549 cells were seeded in a 12-well plate at 1 x 10^5^ cells/well in 1 mL 10% FBS DMEM; simultaneously, cells were infected with rLCMV-EGFP, at an MOI of 0.001. To investigate spread throughout a culture, whole well images were taken at 6 h intervals using an Incucyte Zoom S3 live cell imaging system (Sartorius).

### Immunofluorescence (widefield and confocal microscopy)

Trypsinised A549 cells were seeded onto a 19-mm round glass coverslips (VWR) in a 12-well plate at 1 x 10^5^ cells/well, followed by incubation at 37 °C. At 24 hpi, 1 mL 4% (vol/vol) paraformaldehyde in PBS was added directly on top of the 1 mL infection media for 20 mins at room temperature. After fixation, the cells were washed three times in PBS and then incubated in permeabilisation buffer (0.3% [vol/vol] Triton X-100, 1% [wt/vol] bovine serum albumin [BSA] in PBS) for 10 mins at room temperature. Following permeabilisation, the monolayers were washed three times with PBS and incubated with blocking buffer (1% [wt/vol] BSA in PBS) for 1 h. Subsequently, primary antibody made in the BSA blocking buffer containing NP (1:500, in house [sheep]), FLAG (1:100 [mouse/rabbit]) and HA (1:500 [mouse/rabbit]) was incubated for 1 h at room temperature. The cells were then washed three times with PBS and incubated with corresponding Alexa Fluor 488, 594 and 647 secondary antibodies (Life Technologies; 1:500 in BSA blocking buffer) for 1 h at room temperature in a light protected vessel. Cell monolayers were then washed four times with PBS before addition of appropriate stain/dye; F-actin (Phalloidins, Texas Red, 5 μL/well in 500 μL blocking buffer) and β-tubulin (affimer; 1:200 A gift from Professor Michelle Peckham, University of Leeds, UK), for 1 h in a light protected vessel. Cell monolayers were then washed four times with PBS and mounted onto glass coverslips with the addition of Prolong Gold Antifade reagent with DAPI (Thermo Fisher Scientific), cured, sealed, and stored at 4 °C. Images were then taken on either Zeiss LSM 880 confocal microscopy or Olympus Widefield Deconvolution Microscope and processed using Zen (Blue Edition) software and Fiji (Image J). Line scan analysis was performed utilising Zen (Blue Edition).

### Immunofluorescence (FISH)

FISH probes used were identical to a set previously described [29] and synthesised by Stellaris. The protocol was followed according to manufacturer’s instructions, with probes hybridized overnight at 37 °C. As recommended by manufacturer, FISH probes were visualized using widefield microscopy.

### Immunofluorescence (STED)

Trypsinised A549 cells were seeded onto a 19-mm round glass coverslip (VWR) in a 12-well plate at 1 x 10^5^ cells/well, followed by incubation at 37 °C. At 24 hpi, 1 mL 4% (vol/vol) paraformaldehyde in PBS was added directly on top of the 1 mL infection media for 20 mins at room temperature. After fixation, the cells were washed three times in PBS and the incubated in permeabilisation buffer (0.3% [vol/vol] Triton X-100, 1% [wt/vol] bovine serum albumin [BSA] in PBS) for 10 mins at room temperature. Following permeabilisation, the monolayers were washed three times with PBS and incubated with blocking buffer (1% [wt/vol] BSA in PBS) for 1 h. Subsequently, primary antibody in BSA blocking buffer containing FLAG, (1:100 [rabbit/mouse]) and HA (1:500 [rabbit/mouse]) was incubated for 1 h at room temperature. The cells were then washed three times with PBS and incubated with Fluro 647 secondary antibodies (Abberior STAR red; 1:500 in BSA blocking buffer) for 1 h at room temperature in a light protected vessel. Cell monolayers were then washed four times with PBS before addition of F-actin stain (Phalloidins, Texas Red, 5 μL/well in 500 μL blocking buffer) for 1 h in a light protected vessel. Cell monolayers were then washed four times with PBS and mounted onto glass coverslips with the addition of Prolong Gold Antifade reagent (Thermo Fisher Scientific), cured, sealed, and stored at 4 °C.

### Neutralising antibody assay

Trypsinised A549 cells were seeded in a 12-well plate at 1 x 10^5^ cells/well, followed by incubation at 37 °C for 16-24 h. For pre-treated condition, rLCMV-EGFP (MOI of 0.2) was incubated with 1 mL neutralising antibody (5µg/mL; made in 10% FBS DMEM) for 1 h. Remaining conditions (mock, virus only, virus + antibody [added at 3 hpi]) were subject to incubation with 1 mL 10% FBS DMEM for 1 h. Subsequently, the total 1mL for each condition was added to A549s, and infection allowed to progress. At 3hpi, 5µg/mL neutralising antibody was added inhibiting extracellular viral spread. Total green cell count was measured every 6 h using an Incucyte Zoom S3 live cell imaging system (Sartorius) and normalised for confluency variation. The average of three independent experimental repeats is shown, with error bars showing standard deviation at each time point (n = 3).

## ACKNOWLEDGEMENTS

We acknowledge funding from MRC grant MR/T016159/1 in support of JNB, JF, AS and HNT, BBSRC PhD studentship grant to AS, UKHSA PhD studentship grant to OB. The authors thank Dr Ruth Hughes and Dr Sally Boxall of the bioimaging facility, Faculty of Biological Sciences, University of Leeds, for their expert assistance. We acknowledge the support of Wellcome Trust grant WT104918MA, for provision of the Zeiss LSM880 confocal microscope, the BBRSC grant BB/S019464/1 for the STEDYCon STED microscope, and equipment grant 221538/Z/20/Z from The Wellcome Trust, which supports the use of the Incucyte live cell imaging platform.

## SUPPLEMENTAL FIGURE LEGEND

**Figure S1.**
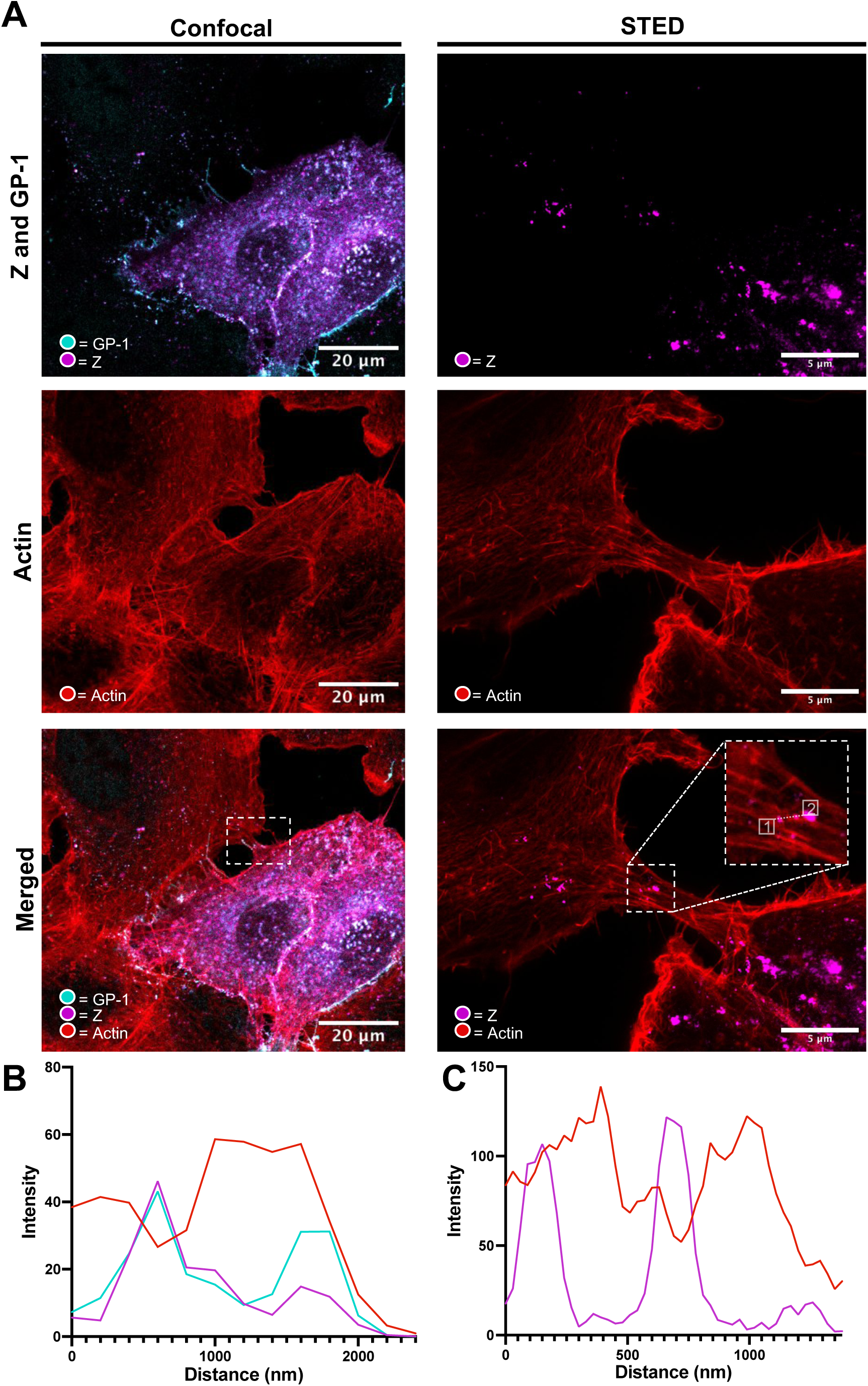
A549 cells infected with rLCMV-GP1-FLAG-Z-HA form actin cell-cell connections where Z and GP-1 closely colocalize. A) A549 cells were infected with rLCMV-GP1-FLAG-Z-HA at an MOI of 0.2. and at 24 hpi, cells were fixed, stained for GP-1-FLAG (cyan), Z-HA (purple) and F-actin (phalloidin, red), then visualized by confocal microscopy (left panels) and stimulated emission depletion microscopy (STED; right panels). (B) Line scan analysis of confocal microscopy image, between points shown on the zoomed image. (C) Line scan analysis of STED, between points shown on the zoomed image.

## REFERENCES

[1] International Committee on Taxonomy of Viruses (ICTV). Taxonomy. 2021 [accessed 2023 May 15]. https://talk.ictvonline.org/taxonomy/.

[2] Centers for Disease Control and Prevention (CDC). Viral Hemorrhagic Fevers. Bunyavirales. [online]. [accessed 2023 May 12]. https://www.cdc.gov/vhf/virus-families/bunyaviridae.htmL

[3] González PH, Cossio PM, Arana R, Maiztegui JI, Laguens RP. Lymphatic tissue in Argentine hemorrhagic fever. Pathologic features. Archives of pathology & laboratory medicine. 1980;104(5):250–4.

[4] Cheng BYH, Nogales A, de la Torre JC, Martínez-Sobrido L. Development of live-attenuated arenavirus vaccines based on codon deoptimization of the viral glycoprotein. Virology. 2017;501:35–46. doi:10.1016/j.virol.2016.11.001

[5] Bonthius DJ, Nichols B, Harb H, Mahoney J, Karacay B. Lymphocytic choriomeningitis virus infection of the developing brain: critical role of host age. Annals of Neurology. 2007;62(4):356–374. doi:10.1002/ana.21193

[6] Bonthius DJ. Lymphocytic Choriomeningitis Virus: An Underrecognized Cause of Neurologic Disease in the Fetus, Child, and Adult. Seminars in Pediatric Neurology. 2012;19(3):89–95. doi:10.1016/j.spen.2012.02.002

[7] Meyer BJ, de la Torre JC, Southern PJ. Arenaviruses: genomic RNAs, transcription, and replication. Current topics in microbiology and immunology. 2002;262:139–57. doi:10.1007/978-3-642-56029-3_6

[8] Maria S. Salvato. Arenaviridae. In: Virus Taxonomy. Elsevier; 2012. p. 715– 723. doi:10.1016/B978-0-12-384684-6.00058-6

[9] Riviere Y, Ahmed R, Southern PJ, Buchmeier MJ, Dutko FJ, Oldstone MB. The S RNA segment of lymphocytic choriomeningitis virus codes for the nucleoprotein and glycoproteins 1 and 2. Journal of virology. 1985;53(3):966–8. doi:10.1128/JVI.53.3.966-968.1985

[10] Byford O, Shaw AB, Tse HN, Todd EJAA, Alvarez-Rodriguez B, Hewson R, Fontana J, Barr JN. Lymphocytic choriomeningitis arenavirus requires cellular COPI and AP-4 complexes for efficient replication and virion production. bioRxiv. 2023 Jan 1:2023.06.27.546563. http://biorxiv.org/content/early/2023/06/27/2023.06.27.546563.abstract. doi:10.1101/2023.06.27.546563

[11] Sakabe S, Witwit H, Khafaji R, Cubitt B, de la Torre JC. Chaperonin TRiC/CCT Participates in Mammarenavirus Multiplication in Human Cells via Interaction with the Viral Nucleoprotein. Journal of Virology. 2023;97(2). doi:10.1128/jvi.01688-22

[12] Bakkers MJG, Moon-Walker A, Herlo R, Brusic V, Stubbs SH, Hastie KM, Saphire EO, Kirchhausen TL, Whelan SPJ. CD164 is a host factor for lymphocytic choriomeningitis virus entry. Proceedings of the National Academy of Sciences of the United States of America. 2022;119(10):e2119676119. doi:10.1073/pnas.2119676119

[13] Pepe A, Pietropaoli S, Vos M, Barba-Spaeth G, Zurzolo C. Tunneling nanotubes provide a route for SARS-CoV-2 spreading. Science advances. 2022;8(29):eabo0171. doi:10.1126/sciadv.abo0171

[14] Aggarwal A, Iemma TL, Shih I, Newsome TP, McAllery S, Cunningham AL, Turville SG. Mobilization of HIV Spread by Diaphanous 2 Dependent Filopodia in Infected Dendritic Cells. PLoS Pathogens. 2012;8(6):e1002762. doi:10.1371/journal.ppat.1002762

[15] Souriant S, Balboa L, Dupont M, Pingris K, Kviatcovsky D, Cougoule C, Lastrucci C, Bah A, Gasser R, Poincloux R, et al. Tuberculosis Exacerbates HIV-1 Infection through IL-10/STAT3-Dependent Tunneling Nanotube Formation in Macrophages. Cell Reports. 2019;26(13):3586–3599.e7. doi:10.1016/j.celrep.2019.02.091

[16] Kadiu I, Gendelman HE. Human Immunodeficiency Virus type 1 Endocytic Trafficking Through Macrophage Bridging Conduits Facilitates Spread of Infection. Journal of Neuroimmune Pharmacology. 2011;6(4):658–675. doi:10.1007/s11481-011-9298-z

[17] Eugenin EA, Gaskill PJ, Berman JW. Tunneling nanotubes (TNT) are induced by HIV-infection of macrophages: a potential mechanism for intercellular HIV trafficking. Cellular immunology. 2009;254(2):142–8. doi:10.1016/j.cellimm.2008.08.005

[18] Martinez MG, Kielian M. Intercellular Extensions Are Induced by the Alphavirus Structural Proteins and Mediate Virus Transmission. PLOS Pathogens. 2016;12(12):e1006061. doi:10.1371/journal.ppat.1006061

[19] Ganti K, Han J, Manicassamy B, Lowen AC. Rab11a mediates cell-cell spread and reassortment of influenza A virus genomes via tunneling nanotubes. PLoS pathogens. 2021;17(9):e1009321. doi:10.1371/journal.ppat.1009321

[20] Mothes W, Sherer NM, Jin J, Zhong P. Virus Cell-to-Cell Transmission. Journal of Virology. 2010;84(17):8360–8368. doi:10.1128/JVI.00443-10

[21] Rustom A, Saffrich R, Markovic I, Walther P, Gerdes H-H. Nanotubular Highways for Intercellular Organelle Transport. Science. 2004;303(5660):1007–1010. doi:10.1126/science.1093133

[22] Shaw AB, Tse HN, Byford O, plahe grace, Moon-Walker A, Saphire EO, Whelan SPJ, Mankouri J, Fontana J, Barr JN. Cellular endosomal potassium ion flux regulates arenavirus uncoating during virus entry. bioRxiv. 2023 Jan 1:2023.06.23.546275. http://biorxiv.org/content/early/2023/06/26/2023.06.23.546275.abstract. doi:10.1101/2023.06.23.546275

[23] Moon-Walker A, Zhang Z, Zyla DS, Buck TK, Li H, Diaz Avalos R, Schendel SL, Hastie KM, Crotty S, Saphire EO. Structural basis for antibody-mediated neutralization of lymphocytic choriomeningitis virus. Cell Chemical Biology. 2023;30(4):403–411.e4. doi:10.1016/j.chembiol.2023.03.005

[24] Jansens RJJ, Tishchenko A, Favoreel HW. Bridging the Gap: Virus Long-Distance Spread via Tunneling Nanotubes. Journal of virology. 2020;94(8). doi:10.1128/JVI.02120-19

[25] Zeng C, Evans JP, King T, Zheng Y-M, Oltz EM, Whelan SPJ, Saif LJ, Peeples ME, Liu S-L. SARS-CoV-2 spreads through cell-to-cell transmission. Proceedings of the National Academy of Sciences. 2022;119(1). doi:10.1073/pnas.2111400119

[26] Kumar A, Kim JH, Ranjan P, Metcalfe MG, Cao W, Mishina M, Gangappa S, Guo Z, Boyden ES, Zaki S, et al. Influenza virus exploits tunneling nanotubes for cell-to-cell spread. Scientific Reports. 2017;7(1):40360. doi:10.1038/srep40360

[27] Lehmann MJ, Sherer NM, Marks CB, Pypaert M, Mothes W. Actin- and myosin-driven movement of viruses along filopodia precedes their entry into cells. Journal of Cell Biology. 2005;170(2):317–325. doi:10.1083/jcb.200503059

[28] Xiao F, Fofana I, Heydmann L, Barth H, Soulier E, Habersetzer F, Doffoël M, Bukh J, Patel AH, Zeisel MB, et al. Hepatitis C Virus Cell-Cell Transmission and Resistance to Direct-Acting Antiviral Agents. PLoS Pathogens. 2014;10(5):e1004128. doi:10.1371/journal.ppat.1004128

[29] King BR, Kellner S, Eisenhauer PL, Bruce EA, Ziegler CM, Zenklusen D, Botten JW. Visualization of the lymphocytic choriomeningitis mammarenavirus (LCMV) genome reveals the early endosome as a possible site for genome replication and viral particle pre-assembly. The Journal of general virology. 2017;98(10):2454–2460. doi:10.1099/jgv.0.000930

